# Multifunctional properties of mycorrhizal helper bacteria: improve mycorrhizal colonization and increase phosphate uptake in Banana

**DOI:** 10.1101/2022.12.13.520192

**Authors:** Chandni Shah, Himanshu Mali, Vijay Kamble, Shachi Mandavgane, R B Subramanian

## Abstract

Using mycorrhizal fungi with their helper bacteria (MHB) to alleviate phosphate deficiency improve plant growth and rejuvenate the soil is the newer safer and most promising environment-friendly approach to limiting synthetic agrochemicals. Here 65 MHBs were isolated from the mycorrhizosphere of the Banana plant and 12 were screened based on their biofilm formation protease and N-acyl homoserine lactones production. A newly reported MHB *Enterobacter* sp. was identified through the 16S ribotyping method and posed plant growth-promoting properties i.e solubilized phosphate produced indole acetic acid ammonia and hydrogen sulfide. The biocompatibility experiment showed a 4-fold increased mycorrhizal proliferation in MHB added plate compared to the control. Further field experiments suggested the banana plants 10-4-fold increment in plant height (216.54 cm) and stem diameter (10.66 cm). At the same time10-fold significant improvements were noted in leaf carbohydrate (639.9 mg g^-1^) protein (432.76 mg g^-1^) chlorophyll (268.66 mg g^-1^) phenol (1.9 mg g^-1^) and proline (uMole g^-1^) content of experimental plant compared to control. The study also showed a 5-fold increase in phosphate content (12.84 mg g^-1^) than the control (2.4 mg g^-1^). The isolates increased the mycorrhizal colonization and spore number by 79.33 % and 14.31 g^-1^ in the rhizospheric soil. Total organic carbon and nitrogen (0.68 %) total phosphorus (55 kg ha^-1^) and potassium (499.66 kg ha^-1^) content of soil were positively affected by MHB and AMF. Further-more principal component analysis (PCA) and Pearsons correlation analysis of all the obtained results clearly showed the positive insights of inoculated MHB and AMF on the growth of the banana plant and soil restoration.

## Introduction

Banana is an agronomically important fruit crop due to their higher nutritional values. India is the leading producer and exporter of bananas worldwide, producing 31.25 million tons (www.statista.com, economictimes.indiatimes.com). Banana cultivation depends on synthetic chemical fertilizers and pesticides (Patel et al., 2018). Current practice increases production costs and causes environmental damage (Mali et al., 2022a). These phosphatic fertilizers are relatively insufficient as they have strong sorption or fixation properties with the soil matrix (Uzma et al., 2022). Ultimately, 80-90% of the soil phosphate (P) is insoluble, immobilized, participates present as a legacy phosphate (P), and cannot be used by plants (Zhu et al., 2018). Soil-dwelling microorganisms have several mechanisms to scavenge the legacy P, mine the soil, and avail P for plants (Zhu et al., 2018).

Microorganism includes bacteria, fungi, algae, protozoans, and nematodes. Amongst those, fungi or mycorrhizal fungi have a symbiotic association with 80% of terrestrial land plants and participate in nutrient transport (Giovannini et al., 2020). They are continuously interlinked with soil bacterial communities and create a “tripartite association” to increase nutrient availability in the soil and plant growth (Giovannini et al., 2020). These symbiotic associations significantly enhance the soil microbial biodiversity and improve soil structure. These symbiotic mycorrhizal helper bacteria (MHB) belong to several genera (*Pseudomonas, Burkholderia, Klebsiella, Enterobacter*, and *Bacillus*) (Reis et al., 2021). The MHBs promote spore germination, stimulate hyphal and mycelium growth, and increase mycorrhizal fungus colonization with the root of host plants (Sundram et al., 2011). Besides, MHBs may be considered plant growth-promoting rhizobacteria (PGPR) (Sangwan and Prasanna, 2021). They can solubilize essential nutrients such as P for plants, synthesize several growth-promoting traits, and release biocontrol agents to protect various abiotic and biotic factors (Bekti et al., 2022).

Thus, elucidating the beneficial effect of this tripartite association between bacteria (MHB)-fungi-plant is an economical, eco-friendly, and sustainable approach to increasing nutrient status, their uptake, and the overall growth and development of a plant (Sangwan and Prasanna, 2021). Here, MHBs act as a PGPR to increase nutrient availability, while AMF and plants work together or individually to benefit plant growth by increasing the absorptions of water and essential minerals nutrients. Moreover, AMF is physically associated with the plant root cortex and increases its hyphal root, ultimately increasing its absorption to absorb the immobile nutrients from the earth’s crust.

In the present study, *Enterobacter cloacae* and *Glomus proliferum* were identified, combined, and co-inoculated on the banana plant to examine their survivability, plant growth (morphological and biochemical characteristics), and leaf phosphate content. Also, the effect of MHB on mycorrhizal colonization and rhizospheric soil was examined by observing root colonization through scanning electron microscopy and physiochemical analysis.

## Materials and methods

### Isolation of mycorrhizal helper bacteria (MHB)

To isolate mycorrhizal helper bacteria (MHB), *Glomus proliferum* mycorrhizal fungus was inoculated in banana plants with garden soil and incubated for five weeks (Shah et al., 2022). Mycorrhizal inoculated plants (mycorrhizosphere) with soil were collected, and for intact spores were collected, the wet sieving and decanting method was followed as described by Pacioni et al. (1991) with minor modification (Pacioni, 1991). For that, 100 g of soil was collected and mixed with 1000 mL of sterile D/W and agitated for 25-30 min at 120 rpm. The resulting suspension was filtered through 710-38 µm filters, and each filtered sieve was dispersed in another sterile D/W and mixed gently with collected spores. The spores were observed under the dissecting microscope at 40× magnification. Single uniform mature spores (slightly yellow, complete with glossy appearance) was collected with a sterile wet needle and inoculated on different growth medium (Luria Bertani, nutrient broth, peptone, king B, and soybean casein agar medium) to isolate mycorrhizal helper bacteria. All the plates were incubated at 30 °C for 5-10 days. Morphologically distinct MHBs were collected, purified, and maintained in Luria Bertani’s medium with 20% glycerol at -20 °C.

### Identification of MHB and their *in vitro* AMF spore germination under monoaxenic conditions

Morphologically district MHBs were preliminarily identified for their morphological, biochemical, and molecular characteristics. Further plant growth-promoting traits and their fungus spore germination assay were performed. As mentioned below, all the MHBs, biofilm formation, protease, and N-AHL production were performed.

### Biofilm formation and motility assay

Biofilm formation was performed using crystal violet as described by Zhao et al. (2021) with minor modification (Zhao et al., 2021). Briefly, exponential phased bacterial cells (A_600_ ≅ 0.05) were inoculated into a 96-well plate and incubated at 30 °C without shaking. After incubation for 24-48 h, the medium was discarded, and cells were washed thrice with Milli-Q water, stained with 0.1 % crystal violet for 20 minutes, and washed with water to remove excess stain and air dry. In the ELISA plate reader, adherent cells that absorb crystal violet were dissolved with 95% acetone and measured at A_570_. At the same time, motility and protease assay of the isolates was performed using Luria Bertani’s medium supplemented with 0.5% agar and skimmed milk agar, respectively. Isolates were spot inoculated (A_600_ ≅ 0.5) at the center of the plates and incubated at 30°C. Swimming motility and zone of hydrolysis were measured, as mentioned by (Zhao et al., 2021).

### Detection of N-AHL production

AHL production was identified using biosensor strain *Chromobacterium violaceum* CV026, which has broad-range detection of short-chain (C2-C8) AHLs (Pérez-Montaño et al., 2013). For identification, a parallel striking method was performed. All isolates were streaked parallel on Luria agar with *C. violence* CV026 here, positive and negative control as *Chromobacterium violence* (MTCC No.2656), and *E-Coil* (MTCC No.443) collected. Plates were incubated at 30°C for 24-48h, and violet coloration indicated the positive.

### *In vitro* mycorrhizal spore germination assay

The MHBs strain was examined to germinate mycorrhizal fungus spores *in vitro* by performing an overlaid monoxenic method using a Modified Strullu and Romand (MSR) medium (Srinivasan et al., 2014) with slight modification. Spores of fungus were spot inoculated at the center, overlaid with 0.5% Luria agar seeded with MHB (10^8^ CFU mL^-1^) and control without MHB, and incubated at 30°C. After 2-4 days of germination, the plates were observed under a stereomicroscope at 40× magnification.

### Morphological, biochemical, and PGP trait characterization

Gram staining and endospore staining were performed as described in Bergey’s manual for microbiology (2001) (Bergey, 2001). Slides were observed under a light microscope at 100× (MPM 400 Eaxeo-PLAN with image analysis system). Biochemical and plant growth-promoting properties such as., Voges-Proskauer (V.P.) test, Methyl Red (M.R.), hydrogen cyanide and hydrogen sulfide (Bakker and Schippers, 1987), indole acetic acid (Uzma et al., 2022), and ammonia production (Iyer et al., 2017), phosphate solubilization (Uzma et al., 2022) and exoenzymes amylase, lipase, urease, catalase, gelatinase production were performed (Dhameliya et al., 2020).

### Molecular identification of the mycorrhizal helper bacteria

Molecular identification was carried out by extracting gDNA, as described by Mali et al. (2022) (Mali et al., 2022a). The 16S rDNA gene sequence was amplified using universal primers 27F and 1492R (forward primer 5’AGA GTT TGA TCC TGG CTC AG 3’ and reverse primer 5’ACG GCT ACC TTG TTA CGA CTT 3’). These PCR conditions were optimized as described by Mali et al. (2022b) (Mali et al., 2022b), as initial denaturation at 95 °C for 3 min followed by denaturation at 95 °C for 30 s, annealing at 51 °C for 30 s and extension 72 °C for 30 s for 25 cycles by final extension 72 °C for 3 min and reaction hold at 4 °C in a thermocycler (Gradient thermocycler® Eppendorf). Amplified gene products were sequenced using the Sanger sequencing method, and sequences were submitted to the NCBI-GenBank database.

### Inoculum preparation and pot experiment

Spore germination assay suggested the bio-compatibility between MHB and mycorrhizal fungus; thus, they were considered for plant growth promotion. For plant inoculation, MHB and mycorrhizal fungus were grown in L.B. and MSR medium 24h at 120rpm. After growth, centrifuged at 5000 ×g for 5 min at R.T., resuspended in sterile D/W, and maintained cell density at 1.5 × 10^7^ CFU mL^-1^ and 7× 10^5^ IP mL^-1^ of MHB and AMF for inoculation on a Banana plant. Four treatments were T1-control, T2-mycorrhizal fungus, T3-MHB, and T4-mycorrhizal fungus + MHB.

Tissue-cultured banana plants seedling (*Musa acuminata AAA*, Cavendish subgroup cultivar William) with 2-3 leaf stages were collected from Vitrigold Biotech Pvt. Ltd, Gujarat, India. Banana plants were uprooted gently, washed twice with sterile distilled water (D/W), and treated per the above treatment. All the plants were potted (pot dimensions are 25 cm long and 30 cm wide) with garden soil (loamy soil with pH 6.5, E.C. (25 °C) 0.91 ds m^-1^, organic carbon 0.36%, total nitrogen 0.031%, available P and K 40.18 kg ha^-1^ and 450.0 kg ha^-1^), with six replicates arranged in completely randomized block design. Field experiments were conducted for 90 days from November to February 2019-2020 at the Botanical Garden of the Department of Biosciences, Sardar Patel University, Gujarat, India. On the 90^th^ day, plants were uprooted gently with their roots and measured for their morphological (plant height, number of leaves, leaf area, and pseudostem diameter) and biochemical parameters (total carbohydrate, total chlorophyll, protein, phosphate, phenol, and proline).

### Biochemical characterization of the pot experiments

For the biochemical analysis, a 2 g leaf sample from all the treated plants was collected and extracted for total carbohydrate (Haisman and Clarke, 1975), total Chlorophyll (Sadasivam and Manickam, 1992), protein (Lowry et al., 1951), phosphate (Soltanpour and Workman, 1979), phenol (Kadmiri et al., 2018), and proline (Patel et al., 2018) was performed.

### Localization of MHB and AMF on banana roots

To identify the localization of MHB and AMF, scanning electron microscopy of treated banana plant roots was collected. Roots were washed twice with sterile D/W, cut into 2-3 mm length segments, and fixed with 2.5% glutaraldehyde in 0.2 M phosphate buffer for 24h (R. Goswami et al., 2018). Selected root samples were dried and mounted on A.I. stubs, coated with gold, and observed under SEM with 5 kV.

### Determination of rhizospheric soil’s nutrient status

All treatments’ rhizospheric soil was collected into sterile autoclaved beg and air-dried. Here, 10 g of soil was analyzed for pH, E.C., total organic carbon, nitrogen, phosphorus, potassium, and other micronutrient assessment at the soil testing laboratory, GSFC, Pvt. Ltd, Gujarat, India.

### Statistical analysis

All the experiments were performed in triplicates, and data were analyzed by performing One-Way ANOVA with Turkey’s Posthoc LSD test (p <0.05) and Pearson’s correlation analysis with principal component analysis using IBM SPSS statistics 22 software. The data were expressed in mean ± standard deviation (S.D.). All the graphs were prepared using a GraphPad prism.

## Results

### Isolation, screening, and identification of MHBs

Morphologically sixty-five distinct MHB strains were isolated from seventeen different spores of the banana rhizosphere. All the isolates were screened for their biofilm formation, protease, and N-acyl homoserine lactones production, as presented in Table 1. Of 65 strains, 27.6% are strong, 36.92% are moderate, and 35.38% are weak producers (Table 1, Figure S1). While 21.53%, 52.32%, and 26.15% are strong, medium, and inefficient protease producers, respectively, 52.30% and 30.76% of isolated strains are motile, and 13.84% are non-motile (Table 1, Figure S2). From the 65 MHB strains, two isolates showed a violet coloration on the *CV026* strain, which indicated the presence of N-AHL production (Table 1, Figure S3).

**Table 1.** Sixty-five MHB isolates primary screening through biofilm formation, protease production, motility and detection of quorum sensing molecules.

| Isolates | Biofilm | Protease | Motility | QS | Isolates | Biofilm | Protease | Motility | QS |
| --- | --- | --- | --- | --- | --- | --- | --- | --- | --- |
| 1 | 2.1592 | + | +++ | - | 34 | 0.6837 | + | ++ | - |
| 2 | 2.3218 | +++ | +++ | - | 35 | 0.9305 | +++ | +++ | - |
| 3 | 3.0759 | + | +++ | - | 36 | 0.829 | +++ | +++ | - |
| 4 | 3.0605 | ++ | +++ | + | 37 | 0.8972 | + | +++ | - |
| 5 | 2.8216 | - | +++ | - | 38 | 0.7205 | + | ++ | - |
| 6 | 1.8264 | - | +++ | - | 39 | 0.8231 | - | ++ | - |
| 7 | 3.076 | + | ++ | - | 40 | 0.86 | - | +++ | - |
| 8 | 3.3533 | - | +++ | - | 41 | 1.63 | + | +++ | - |
| 9 | 3.1304 | - | +++ | - | 42 | 1.8352 | + | ++ | - |
| 10 | 3.1003 | + | ++ | - | 43 | 1.3523 | + | ++ | - |
| 11 | 0.646 | - | ++ | - | 44 | 0.8811 | + | +++ | - |
| 12 | 0.647 | ++ | +++ | - | 45 | 0.6944 | - | ++ | - |
| 13 | 1.158 | +++ | +++ | - | 46 | 2.7918 | +++ | ++ | - |
| 14 | 1.9522 | +++ | +++ | - | 47 | 0.6944 | ++ | ++ | - |
| 15 | 1.3959 | - | +++ | - | 48 | 3.0032 | - | ++ | - |
| 16 | 0.7816 | +++ | +++ | - | 49 | 2.3533 | + | ++ | - |
| 17 | 0.9012 | - | +++ | - | 50 | 1.3413 | +++ | ++ | - |
| 18 | 1.0522 | + | +++ | - | 51 | 2.6987 | + | +++ | - |
| 19 | 0.7771 | - | +++ | - | 52 | 3.2041 | - | + | - |
| 20 | 1.6636 | ++ | ++ | - | 53 | 2.8988 | - | + | - |
| 21 | 1.7606 | + | +++ | - | 54 | 2.8416 | - | + | - |
| 22 | 1.5818 | - | +++ | - | 55 | 1.6433 | + | ++ | - |
| 23 | 0.6997 | + | +++ | - | 56 | 2.7617 | ++ | +++ | - |
| 24 | 0.6859 | - | +++ | - | 57 | 1.2687 | - | + | - |
| 25 | 1.2852 | - | +++ | - | 58 | 2.8255 | ++ | + | - |
| 26 | 1.1012 | +++ | +++ | - | 59 | 0.7442 | - | + | - |
| 27 | 0.688 | +++ | +++ | - | 60 | 2.8699 | - | + | - |
| 28 | 1.0458 | +++ | +++ | - | 61 | 1.8595 | + | + | - |
| 29 | 1.2474 | +++ | +++ | - | 62 | 0.7986 | - | ++ | - |
| 30 | 1.4138 | +++ | +++ | - | 63 | 1.3393 | - | - | - |
| 31 | 0.7788 | - | ++ | - | 64 | 0.067 | - | ++ | - |
| 32 | 1.004 | ++ | +++ | - | 65 | 0.2022 | +++ | + | + |
| 33 | 1.0932 | + | ++ | - | (Strong = +++, moderate = ++, weak = +, and not observed = -) |  |  |  |  |

While these two isolates (MHB 4 and 65) have the most potential properties, the MHB4 strain was again characterized for *In Vitro* mycorrhizal fungus spore germination through overlaid monoxenic plate method. From the microscopic observations, the MHB4 inoculated spores germinated earlier and showed more dense spores and hyphal network growth than the control in Figure 1. The results also suggested that both the MHB strain and mycorrhizal fungus have biocompatibility by observing both organisms’ increased biomass. Among the 65 isolates, the twelve most potent isolates were further screened for plant growth-promoting, biochemical, morphological, and molecular identification.

**Figure 1.**
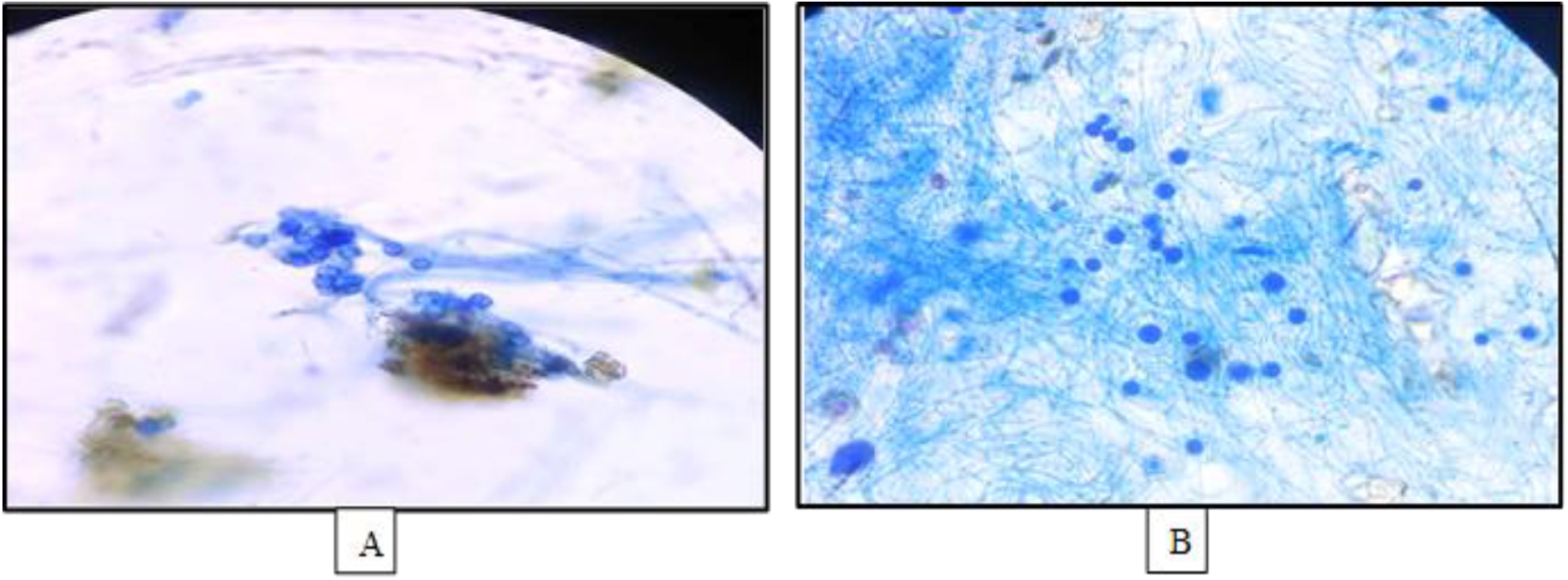
AMF spore germination assay was performed with MHB4CP under monoxenic MSR agar medium and observed at a 40 X microscope. A) AMF & B) MHB4CP +AMF.

### Identification of MHB

#### Morphological identification and Biochemical and PGPR characterization

Twelve putative MHB strains were purified from the banana rhizospheric fungus spores, and all were morphologically identified by gram staining and endospore straining. Of the 12, eight MHB strains were identified as gram-negative, small, and medium rods. At the same time, three are gram-positive, medium rods, and one belongs to gram-negative, spherical cocci in nature, as presented in Table 2 and Figure S4. Most are white or slightly yellow-colored, wet, and opaque in physical appearance. All the isolates are endospore-forming except one MHB2. All the isolates appear favorable for the methyl red and Voges-Proskauer test. Also, release several exoenzymes such as amylase, lipase, urease, catalase, and gelatinase.

**Table 2.**
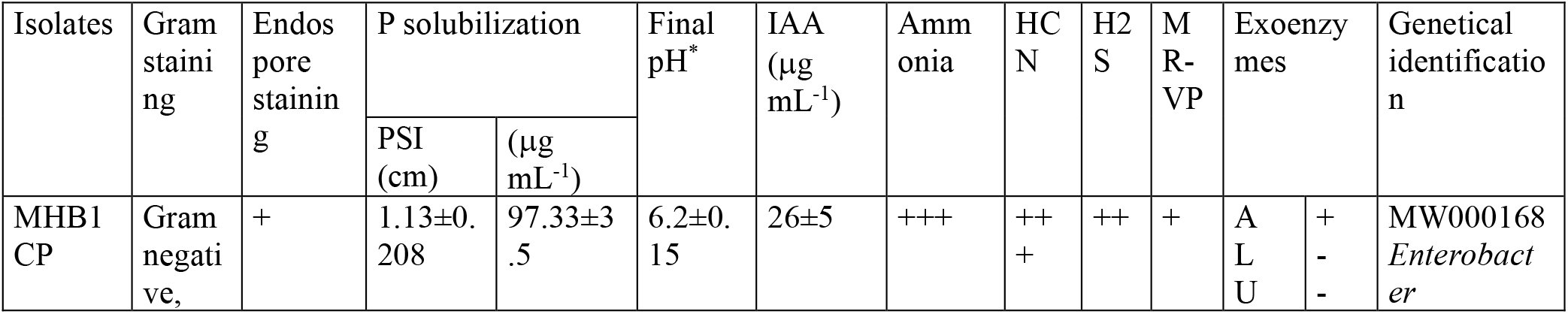
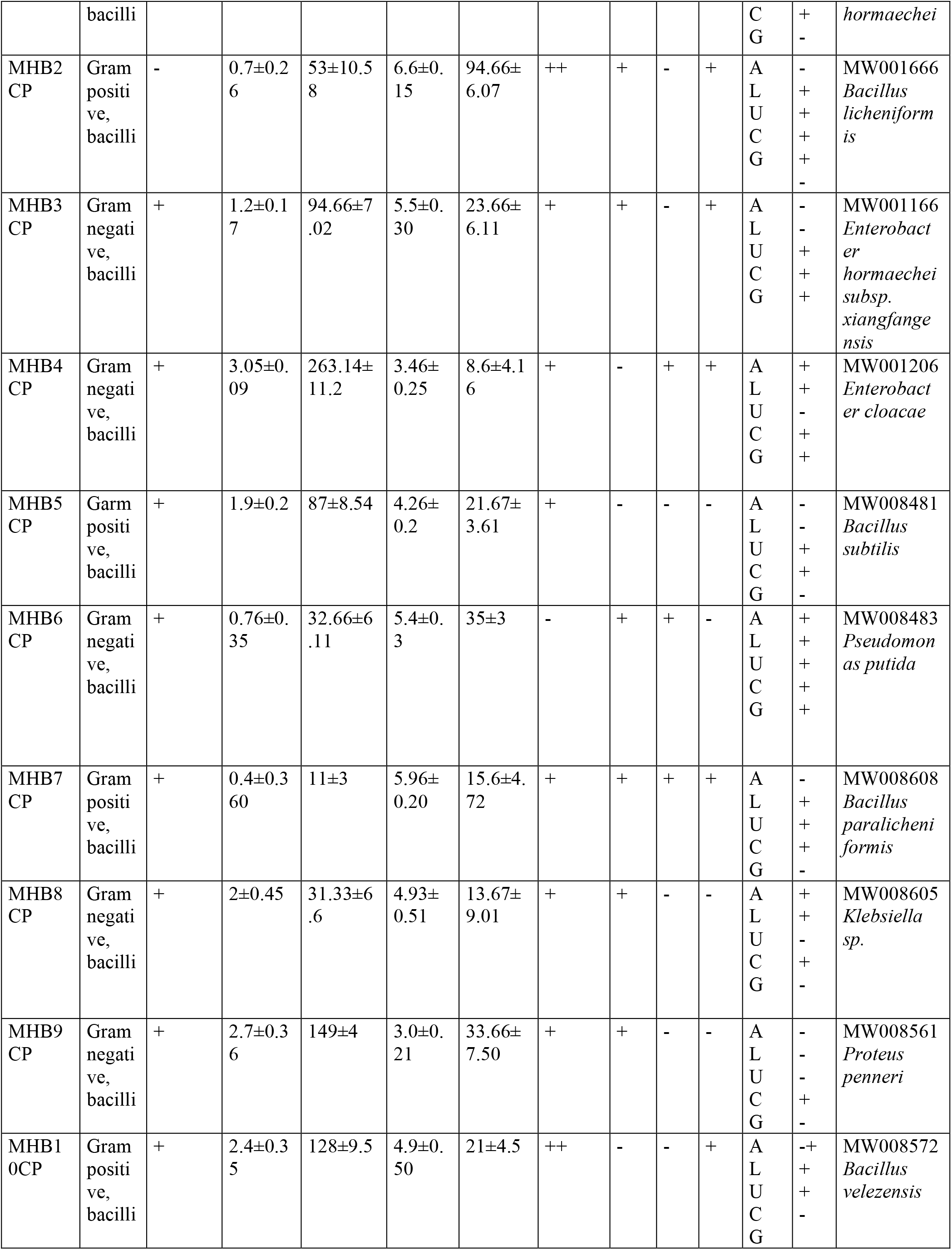
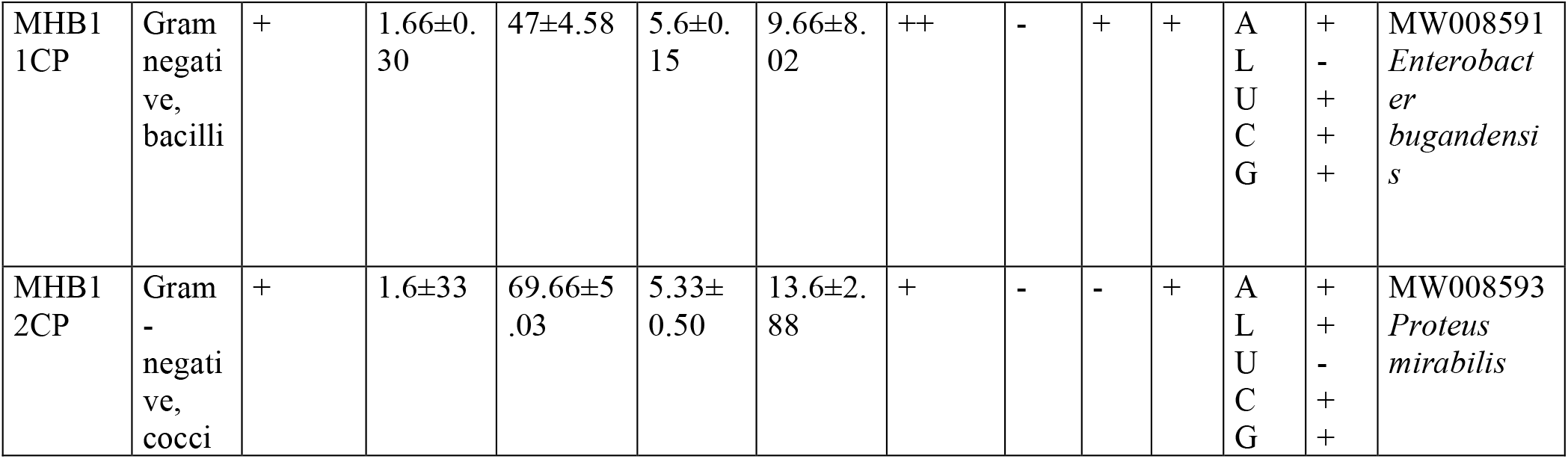
MHB isolate morphological identification, PGPR characteristics such as P solubilization, Indole acetic acid, ammonia, hydrogen cyanide, hydrogen sulfide, MR-VP, and exoenzymes production, with molecular identification with GenBank accession number. All the results were expressed as mean ± standard deviation (SD). Here, PSI, IAA, HCN, H_2_S, MR-VP, A, L, U, C, and G are Phosphate solubilization index, Indole acetic acid, hydrogen cyanide, hydrogen sulfide, methyl red-Voges-Proskauer, amylase, lipase, urease, catalase, and gelatinase, respectively. Tricalcium phosphate were used as inorganic insoluble source of P for P solubilization. Here Strong = +++, moderate = ++, weak = +, and not observed = -.

Based on plant growth-promoting traits, all the isolated strains solubilized insoluble phosphate by measuring the phosphate solubilization index (PSI), releasing soluble phosphate (P), and declining the medium’s pH. Results suggested that all the isolates showed PSI in the range from 0.4 to 3.05 cm, while soluble P ranged from 11 to 263.11 µg mL^-1^ and declined from pH 3.03 to 6.6 from initial final pH 7, as presented in Table 2. Next to P solubilization, all the isolates produced indole acetic acid and ammonia, two significant plant growth-promoting components. All the isolates produced IAA concentrations between 8.6 to 94.66 µg mL^-1^. Most isolates showed the presence of hydrogen cyanide and sulfide (Table 2). However, identifying the bacterial isolates through morphological, biochemical, and PGPR characteristics is difficult; thus, all the isolates were molecularly identified through 16S rRNA gene sequencing. From the NCBI GenBank database, isolates belong to the genera *Enterobacter, Bacillus, Proteus, Pseudomonas*, and *Klebsiella*. In particular, MHB (1, 3, 4, and 11) is classified into *Enterobacter hormaechei and Enterobacter hormaechei subsp. xiangfangensis, Enterobacter cloacae*, and *Enterobacter bugandensis*, while MHB (2, 5, 7, and 10) identified as *Bacillus licheniformis, Bacillus subtilis, Bacillus paralicheniformis*, and *Bacillus velezensis*, respectively. The other MHBs (6, 8, 9, and 12) were *Pseudomonas putida, Klebsiella sp*., *Proteus penneri*, and *Proteus mirabilis*. All the isolates 16S gene sequences were submitted to the NCBI GenBank, and their accession numbers are MW000168, MW001666, MW001166, MW001206, MW008481, MW008483, MW008608, MW008605, MW008561, MW008572, MW008591, and MW008593 as present in Table 2.

From the results of primary screening, biochemical, and PGPR characteristics, isolated strain MHB4CP showed the highest phosphate solubilization, poses all the other properties, and biocompatibility with mycorrhizal fungus were further considered for their plant growth promotion activity on Banana to alleviate the P deficiency and revives the soil.

### Growth-stimulatory effect of MHB and AMF on Banana at the field level

Inoculation of the isolates significantly enhanced the plant height, pseudostem diameter, and no of leaves, as presented in Table 3. The highest increment was observed in the MHB and AMF (T4) inoculated plants, followed by MHB (T3) and AMF (T2) compared to the control (T1). In particular, T4 showed a maximum increment of plant height by recording 216.65 from an initial 14.2 cm, pseudostem diameter 10.66 to 1.53 cm, and no leaves from 12.33 to 2. The post-plant study also revealed the same and showed that shoot and root fresh and dry weight were also positively affected by the inoculation of isolates. Results mentioned that T4 led the highest importance, 583.02 (SFW), 34.57 (SDW), 688.866 (RFW), and 52.39 (RDW) compared to T3, T2, and T1, as shown in Table 3.

**Table 3.** Plant growth parameters of various treatment (T1-T4) on bananas treated with MHB, AMF, and combination. The mean of plant height, shoot fresh weight (SFW), shoot dry weight (SDW), root fresh weight (RFW), root dry weight (RDW), pseudostem diameter, and the number of leaves were evaluated on day-90 after inoculation into pot experiments. Different superscripted alphabets over the standard deviation in a column represent the significant difference of One-way ANOVA followed by Turkey’s Posthoc LSD test p < 0.05. (Number of replicates = 6)

| Treatments | Plant physiological parameters |  |  |  |  |  |  |
| --- | --- | --- | --- | --- | --- | --- | --- |
|  | Height (cm) | SFW (g) | SDW (g) | RFW (g) | RDW (g) | Pseudostem diameter (cm) | Number of leaves |
| T1 | 21.38±6.30<sup>d</sup> | 126.54±10.7<sub>d</sub> | 7.1±1.49<sup>cd</sup> | 120.72±7.14<sub>d</sub> | 20.41±4.02<sub>cd</sub> | 2.53±1.02<sup>cd</sup> | 4.33±1.52<sup>cd</sup> |
| T2 | 148.62±17.83<sub>bc</sub> | 428.04±20.9<sub>c</sub> | 26.88±6.55<sub>ab</sub> | 463.70±7.23<sub>b</sub> | 33.38±4.13<sub>bc</sub> | 6±1.37<sup>bc</sup> | 8.6±0.69<sup>bc</sup> |
| T3 | 174.62±9.41<sup>b</sup> | 525.61±15.0<sub>2<sup>b</sup></sub> | 16.48±3.65<sub>bc</sub> | 402.37±8.39<sub>c</sub> | 42.82±8.59<sub>ab</sub> | 7.60±1.13<sup>ab</sup> | 8±2.64<sup>ab</sup> |
| T4 | 216.65±23.02<sub>a</sub> | 583.02±21.2<sub>7<sup>a</sup></sub> | 34.57±4.51<sub>a</sub> | 688.86±17.6<sub>6<sup>a</sup></sub> | 52.39±10.0<sub>1<sup>a</sup></sub> | 10.66±1.82<sub>2<sup>a</sup></sub> | 12.33±3.51<sub>1<sup>a</sup></sub> |

### Effect of inoculation on leaf nutrient content

The leaf nutrients content analysis also finds similarities with the results of morphological parameters and observed that T4 is the most effective by increasing the content of carbohydrates from an initial 55.67 to 652.32 mg g^-1^, chlorophyll from 41.29 to 426.57 mg g^-1^, protein from 23.41 to 267.45 mg g^-1^, phosphate from 2.14 to 14.57 mg g^-1^, proline from 0.41 to 1.67 µMole g^-1^and phenol from 0.78 to 1.94 mg g^-1^ compared to T3, T2, and T1 as control (Figure 2 A, B & C). Phosphate content is improved in AMF and MHB inoculated plants than in control. Overall, the nutrient content of leaves in all the inoculated treatments was significantly 10 to 6-fold higher and improved.

**Figure 2.**
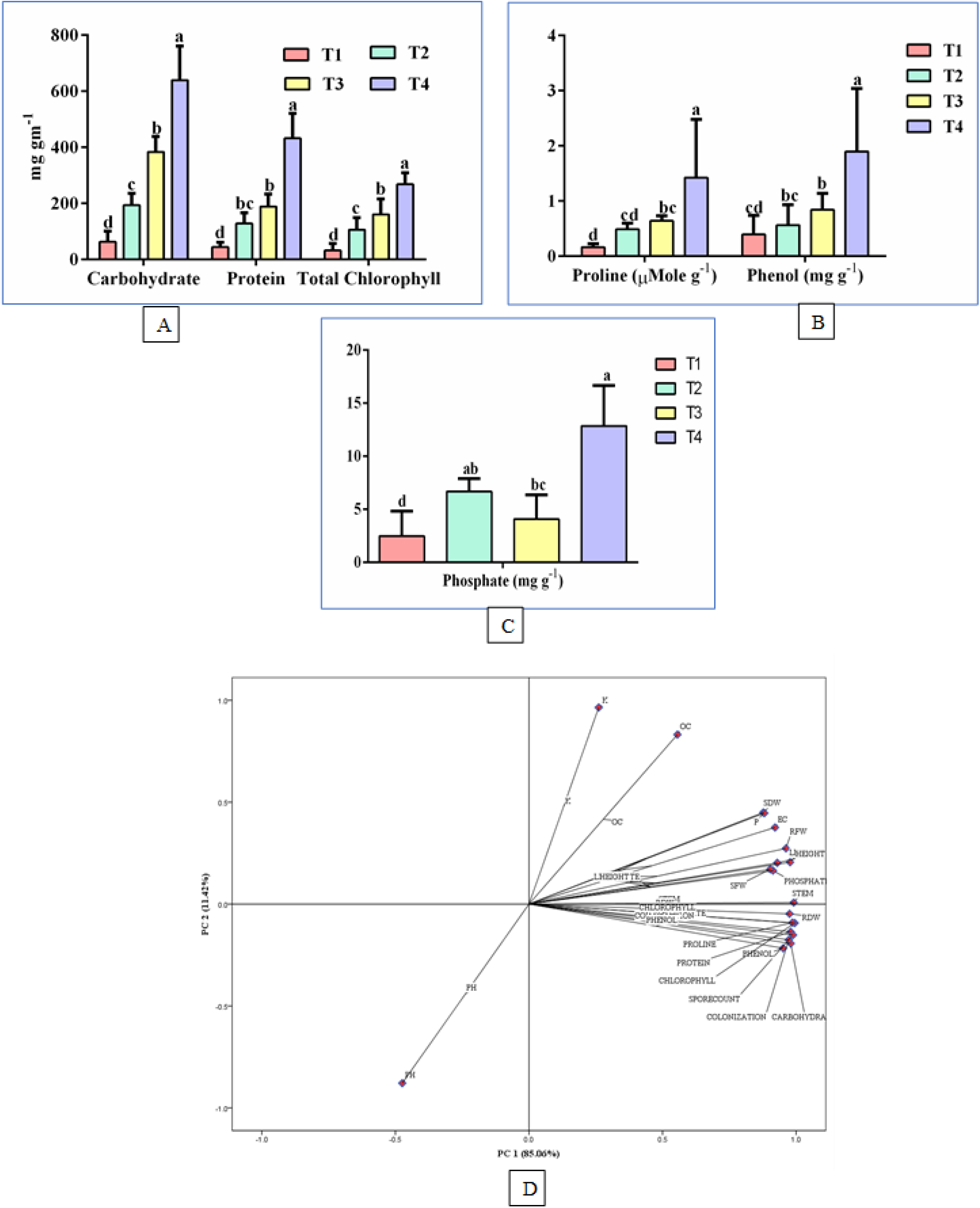
Growth stimulatory effect on various treatments (T1-T4) on biochemical parameters (A) carbohydrate, protein and Chlorophyll, (B) Proline and total phenol content (C) Phosphate of the banana plant under pot experiment and (D) PCA analysis of all the plant and soil parameters. Different superscripted alphabets over the standard deviation in a column represent the significant difference of One-way ANOVA followed by Turkey’s Posthoc LSD test p < 0.05. (Number of replicates = 6)

### Localization of MHB strain on banana roots

The inoculated strains’ localization was identified by scanning electron microscopy of the root on the 90^th^ day after inoculation. The SEM imaging of all the treatments identified the presence of mycorrhizal fungus and MHB. Several bacterial cells and mycorrhizal fungus spore populations were observed, while on the other side, control roots did not find any bacteria or fungi, as observed in Figure 3. Moreover, a mycorrhizal colonization study showed that T4 showed 79.33% and 14.31 g^-1^ colonization and spore accumulation, respectively. From the results, the T3 (MHB) inoculated plant also observed 57.66% and 9.7 g^-1^ colonization and volume of spore compared to the T2 (AMF) inoculated plant with 34.33% and 7.33 g^-1^, suggesting the positive influence of inoculated bacteria on soil’s other mycorrhiza fungus as present in Table 4.

**Table 4.** physicochemical properties of the experimental rhizospheric soil. P-Phosphate, OC- a ratio of organic carbon & nitrogen and K-Potassium. Different superscripted alphabets over the standard deviation in a column represent the significant difference of One-way ANOVA followed by Turkey’s Posthoc LSD test p < 0.05. (Number of replicates = 3)

| Treatments | Soil physicochemical parameters |  |  |  |  |  |  |
| --- | --- | --- | --- | --- | --- | --- | --- |
|  | pH | EC (mS cm<sup>-1</sup>) | P (kg ha<sup>-1</sup>) | %OC | K (kg ha<sup>-1</sup>) | Mycorrhizal colonization (%) | Spore count (g<sup>-1</sup> soil) |
| T1 | 7.33±0.15<sup>a</sup> | 0.36±0.04<sup>d</sup> | 8.51±1.33<sup>d</sup> | 0.37±0.05<sup>cd</sup> | 359.33±5.50<sup>d</sup> | 17.33±5.03 | 5.03±0.65 |
| T2 | 6.20±0.18<sup>b</sup> | 0.54±0.03<sup>ab</sup> | 25±3<sup>c</sup> | 0.53±0.07<sup>bc</sup> | 482.33±4.72<sup>c</sup> | 34.33±4.04 | 7.33±0.90 |
| T3 | 6.93±0.06<sup>cd</sup> | 0.47±0.03<sup>cd</sup> | 42±3<sup>b</sup> | 0.81±0.06<sup>d</sup> | 455.66±12.89<sup>a</sup> | 57.66±3.05 | 9.7±0.8 |
| T4 | 6.63±0.122<sup>c</sup> | 0.633±0.02<sup>a</sup> | 55±4<sup>a</sup> | 0.68±0.08<sup>ab</sup> | 499.66±9.07<sup>bc</sup> | 79.33±6.02 | 14.31±.053 |

### Effect of inoculation on rhizospheric soil

Microorganisms MHB and AMF significantly improved the nutrient content of rhizospheric soil compared to the control (Table 4). Detailed soil analysis found that soil phosphate, potassium, carbon, and nitrogen content were more positively influenced in T4 than in other treatments. Interestingly, T3 (MHB) inoculated soil showed a high K, P, C, and N 455.66 kg ha^-1^, 42 ka ha^-1,^ and 0.81%, respectively. Lastly, the correlation analysis of all the plant morphological, biochemical, and soil physicochemical parameters was performed using SPSS. From the correlation analysis, all the obtained results were found to be positively significant except the pH of the soil (Figure 2D). Here, PCA analysis also considers similarities with the results of correlation analysis, and 95% of the results are directed toward the positive quadrant, suggesting a strong positive correlation between all the considered parameters (Figure 2D). Here, also effects of pH diverged at a robust negative correlation (Table S1).

## Discussion

Several genera of MHB, such as *Pseudomonas, Agrobacterium, Azospirillum, Bacillus, Enterobacter*, and *Burkholderia*, showed a significant potential to propagate mycorrhizal fungi (Sangwan and Prasanna, 2021). These isolates are also considered biofertilizers to increase nutrient availability (specifically P, K, and N) for plants and rejuvenate soil through various cellular and metabolic mechanisms (Frey-Klett et al., 2007). Isolation and screening of the MHB strains from the rhizospheric habitat are difficult, as most of the fungi and their fruiting bodies reside deeper from the plant roots. Thus, the mycorrhizosphere was manipulated by inoculating fungus spores on tissue culture plants. Reis et al. (2021) and Zhao et al. (2014) also demonstrated the identification of MHBs from the mycorrhizosphere of ectomycorrhizal fungus *Pisolithus albus* and *Lactarius rufus*, respectively (Zhao et al., 2014; Reis et al., 2021). However, few literatures mentioned that mycorrhizal association at the root is not necessary for identifying MHB (Labbé et al., 2014; Zhao et al., 2014). One of the reports on poplar plants revealed that bacteria isolated from the phyllo-sphere of the plant could also propagate the mycorrhizae (Zhao et al., 2014). These observations also indicate that MHB isolates are abundant in nature.

In the present study, a wide range of MHB belongs to *Enterobacter, Bacillus, Pseudomonas, Klebsiella*, and *Proteus*, but *Enterobacter* is the most effective mycorrhizal stimulatory bacterium (Table 2). Isolate *Enterobacter* sp. MHB4CP strain increases total mycorrhizal colonization and spore count by 79.33% and 14.31 g^-1^ compared to control by 17.33% and 5.03 g^-1^ from the field study (Table 4). Best of our literature, this is the first report to explain bacterium’s role in mycorrhizal spore germination and its proliferation in soil. We explained that isolated strain MHB4CP is a biofilm-forming, gram-negative, spore-forming, rod-shaped motile bacterium that can solubilize insoluble inorganic phosphate, decline pH, produce IAA, ammonia, hydrogen sulfide, N-Acyl homoserine lactones and released enzymes such as amylase, lipase, catalase, and gelatinase (Table 1 and 2). Previous reports also identified *Enterobacter* sp., which has a growth stimulatory effect on mycorrhizal colonization, improving plant and soil growth and nutrient uptake (Ravn et al., 2001; Khalifa et al., 2016; Pérez-Rodriguez et al., 2020).

Previous studies explained that MHBs play a significant role in mycorrhiza’s growth stimulation and colonization, but the exact mechanism between these interactions is still unknown (Rigamonte et al., 2010; Labbé et al., 2014). Literature also supported that MHB help increases the colonization by increasing the biomass of fungus but did not show an increment in spore accumulation (Labbé et al., 2014; Zhao et al., 2014). Our study also finds a similar observation, where colonization is significantly enhanced, but spore accumulation is limited; however, spore accumulation is high in T4 compared to control (T1) and AMF (T2) alone (Table 4). Similar results were obtained by Reis et al. (2021) and Sundram et al. (2011), where MHB strains *Bacillus* and *Pseudomonas* promote fungus growth but limit spore formation (Sundram et al., 2011; Reis et al., 2021). Reports explained that various unknown volatile organic compounds, known lactone compounds, and some secondary metabolites play a vital role in the interaction between bacterium and fungi (Kurth et al., 2013; Pérez-Montaño et al., 2013). Moreover, these compounds are also responsible for growth stimulation and mycorrhizal germination by binding at membrane steroid binding protein-1, also known as the MYC factor (Boyno and Demir, 2022). The MYC factor activates several plant genes responsible for starch accumulation, calcium spiking, and lateral root elongation (Das et al., 2022).

Literature proposed that MHBs mostly colonized and proliferated at the hyphal and spore surface to influence mycorrhizal growth (Labbé et al., 2014). The present study also finds the same from the localization analysis, where a large population MHBs was observed on the spore surface (Figure 3 T4). In-depth analysis by scanning electron microscopy also identifies the same attachment site where helper bacteria proliferate. Again, isolate MHB4CP and AMF was implemented for their banana plant growth promotion to unrevealed the plant growth-promoting properties of these organisms can prove to be beneficial in alleviating the phosphate deficiency. Results of the field study showed that both AMF and MHB4CP inoculation promote the growth of the banana plant in terms of increasing the plant height, the number of leaves and the pseudostem diameter of the plant compared to the control (Table 3). This event may happen due to the solubilization of minerals and the production of phytohormones and compounds, which may solubilize phosphate and increase availability. A similar observation was reported by Reis et al. (2021) and Sangwan et al. (2021), where inoculated organisms significantly improved plant height and leaf biomass (Reis et al., 2021; Sangwan and Prasanna, 2021).

The nutrient analysis of the leaf also finds the increment in the carbohydrate, total chlorophyll, protein, phosphate, proline, and phenol content of the T4 plant, followed by T2 and T3 than control (Figure 2). This observation finds consistency with Bourles et al. (2020), Pathak et al. (2019), and Wahid et al. (2020), where plants showed high content of total chlorophyll, protein, and phosphate in cotton grass, wheat, and potato (Pathak et al., 2019; Bourles et al., 2020; Wahid et al., 2020). The improvement in nutrient content may be due to ammonia production or the release of exoenzymes that convert complex minerals and nutrients into plant-absorbable form (Ma et al., 2016). Mycorrhizal fungi are responsible for nutrient uptake and transport. Moreover, they are non-symbiotic N-fixers, indirectly increasing N-fixation (Giovannini et al., 2020). Fungi absorbed, scavenged, and mobilized the phosphate and potassium through their well-developed hyphal network from spatially presented P (Giovannini et al., 2020). MHB and mycorrhizal fungi combinedly translocated, mineralized, and mobilized the mineral nutrient to the plant (Sundram et al., 2011). Also, MHB acts as a positive influencer to interact with mycorrhizal fungi to increase colonization. This MHB-fungi interaction fixed the nitrogen, mineralized the phosphate and potassium, and released phytohormones. A significant increment in the plant leaves nutrient content was reported earlier when MHB and mycorrhizal fungus were combined and inoculated on potatoes, wheat, and maize (Bharadwaj, 2007; Liang et al., 2018). Reports also suggested that fungi increase the surface area through hyphal biomass to increase nutrient acquisition from the root and the rhizospheric regions (Deveau et al., 2007; Sharma et al., 2020).

Rhizospheric soil nitrogen, carbon, phosphorus, and potassium contents are considerably enhanced in the treated plant compared to the control (Table 4). This improvement suggested an essential soil fertility factor to increase the accumulation and transfer of nutrients in soil and plants. Nutrient content increased through biochemical processes affected by MHB and AMF metabolic activity performed in the soil (Frey-Klett et al., 2007). These organisms released various enzymes, organic acids, and metabolites to decrease the pH, increasing the soil’s conductivity (Giovannini et al., 2020; Wahid et al., 2020). Giovannini et al. (2020) and Labbe et al. (2014) also showed the response of bacterium and AMF and showed a two-to-four-fold increment in soil nutrient content (Labbé et al., 2014; Giovannini et al., 2020). This study also reported increased rhizospheric soil nutrient quantities after being treated with microorganisms.

The MHB has the biofilm formation ability to be trapped at mycorrhizal fungus hyphae and interact through N-AHL molecules, which is the preliminary observation of the proposed study. Secondly, MHB reacts as a plant growth-promoting rhizobacteria to improve plant growth by providing mineral nutrients and benefits in fungus proliferation to increase the hyphal network at both the root and rhizospheric zone. Plant and soil analysis revealed a positive significant improvement and accumulation of nutrients after being treated with MHB and mycorrhizal fungus. The localization study found a large proportion of bacterial proliferation at the mycorrhizal fungus’s hyphal and spore surface symbiotically associated with plant roots. A further field study has been initiated to identify the mechanisms between MHB and mycorrhizal fungus interaction to identify their positive consequences for alleviating phosphate deficiency, improving plant growth and development, and rejuvenating the soil sustainably.

## Acknowledgment

The author would like to thank Prof. Anuradha Nerurkar and Mr. Ashtaad Vesuma from M. S. University, Baroda, for providing *C. violaceum CV026* (AHL-Bioreporter) strain.

The authors have declared that no funds/grants or other support were received during the preparation of this manuscript. Author C.S. is thankful to ScHeme of Developing High-quality research (SHODH), Government of Gujarat, for financial aid, and author H.M. is grateful to the CSIR-NET-JRF/SRF fellowship for financial support.

## Authors contribution

RBS: Conceptualization, supervision, and resources C.S.: Methodology and writing original draft, C.S. and H.M.: Formal Data analysis and investigation, C.S., V.K., and S.M.: experimentation and data curation, C.S. and RBS: reviewing, editing, and finalizing

## Supplementary file

**Figure S1.**
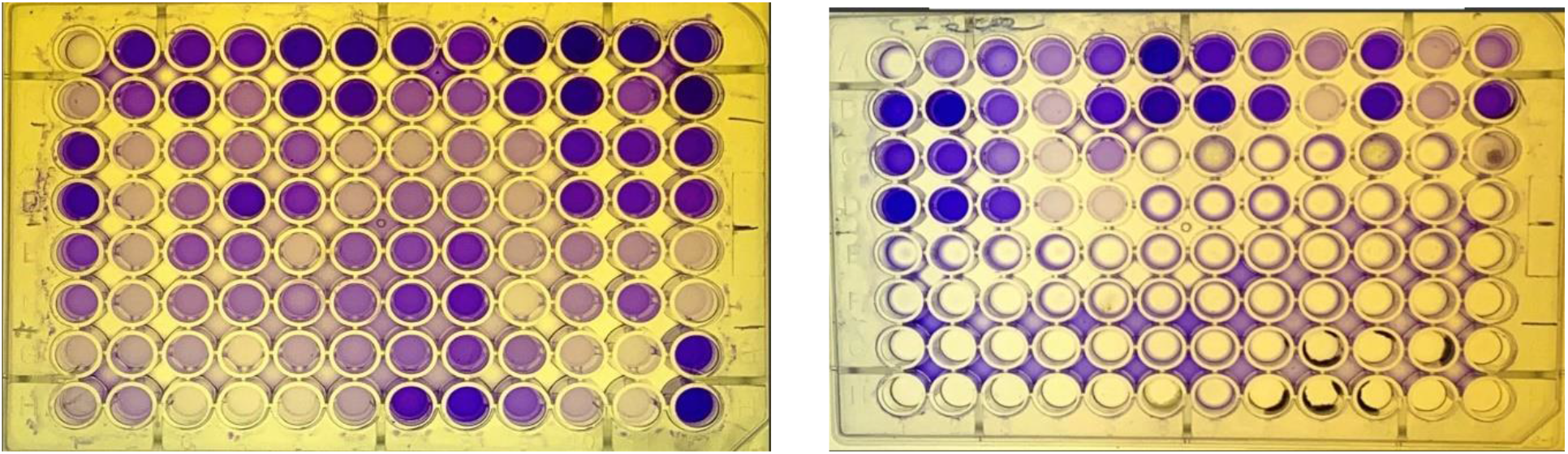
shows the Biofilm formation by MHB isolates.

**Figure S2.**
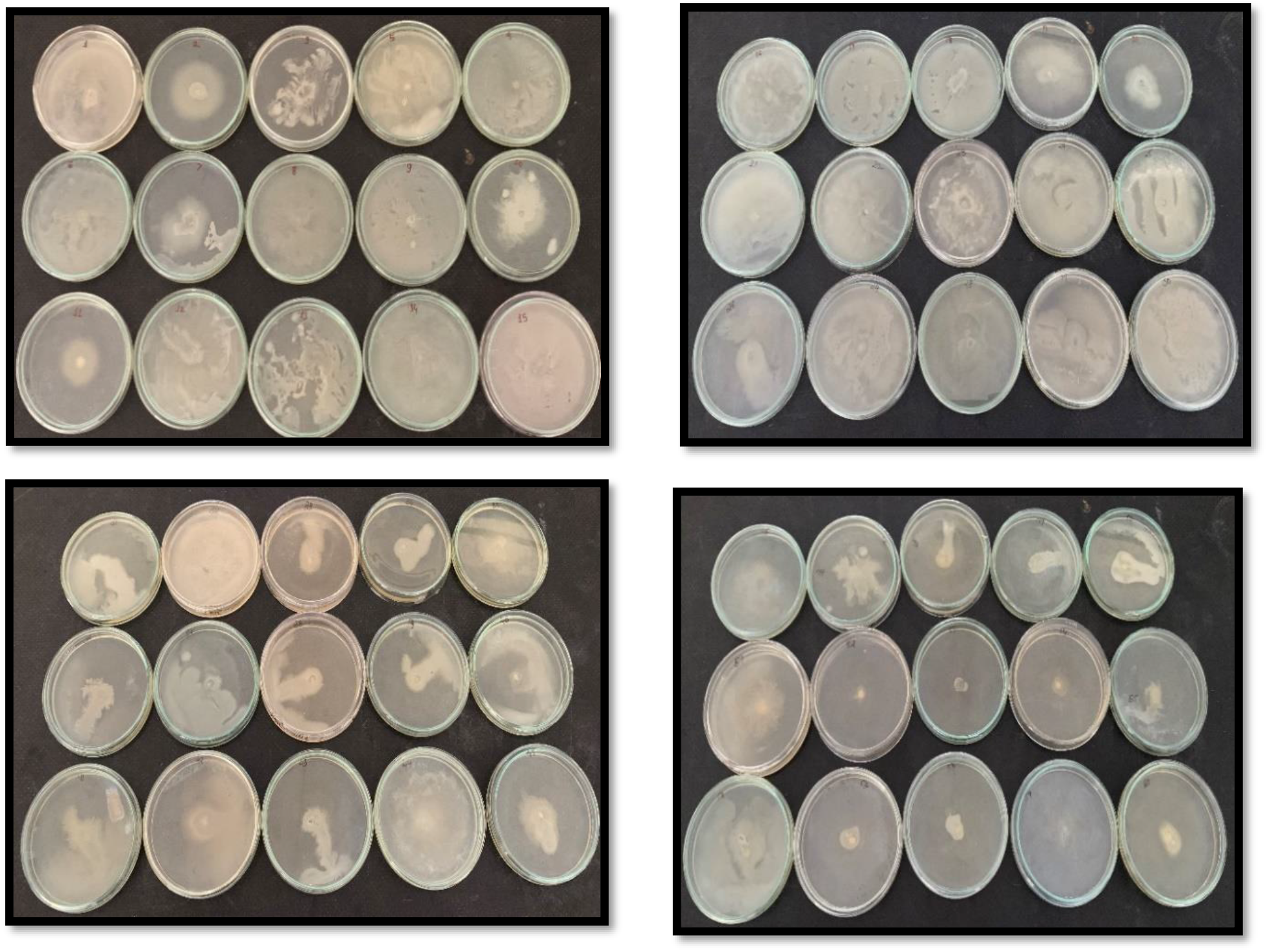
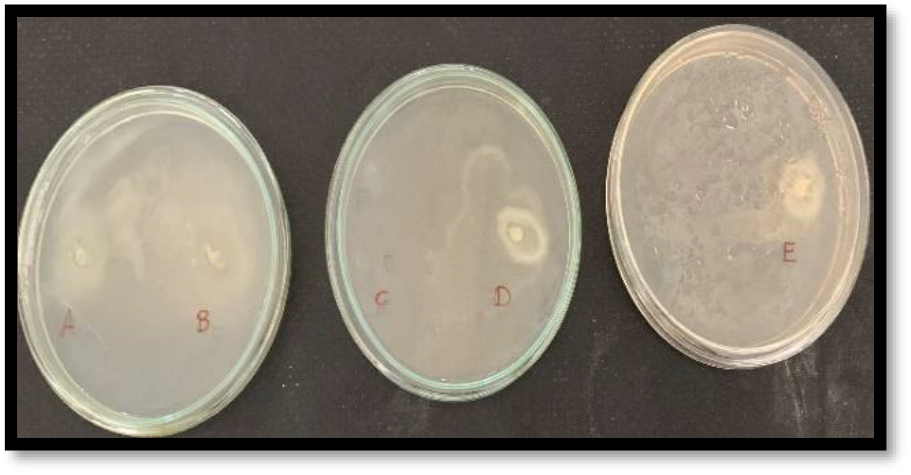
Analysis of swimming motility of MHB isolates

**Figure S3.**
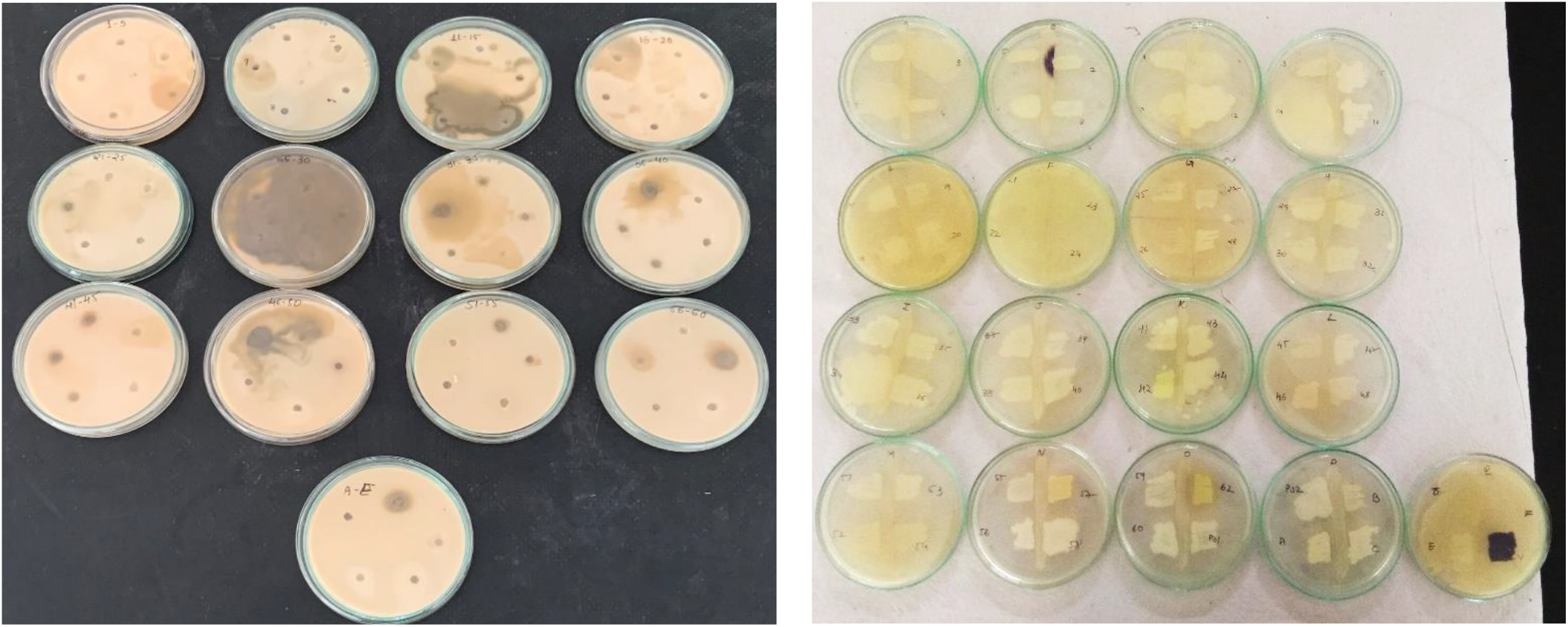
MHB isolates showed protease hydrolysis and AHL production (clear zone of hydrolysis and violet coloration, characterized as positive)

**Figure S4.**
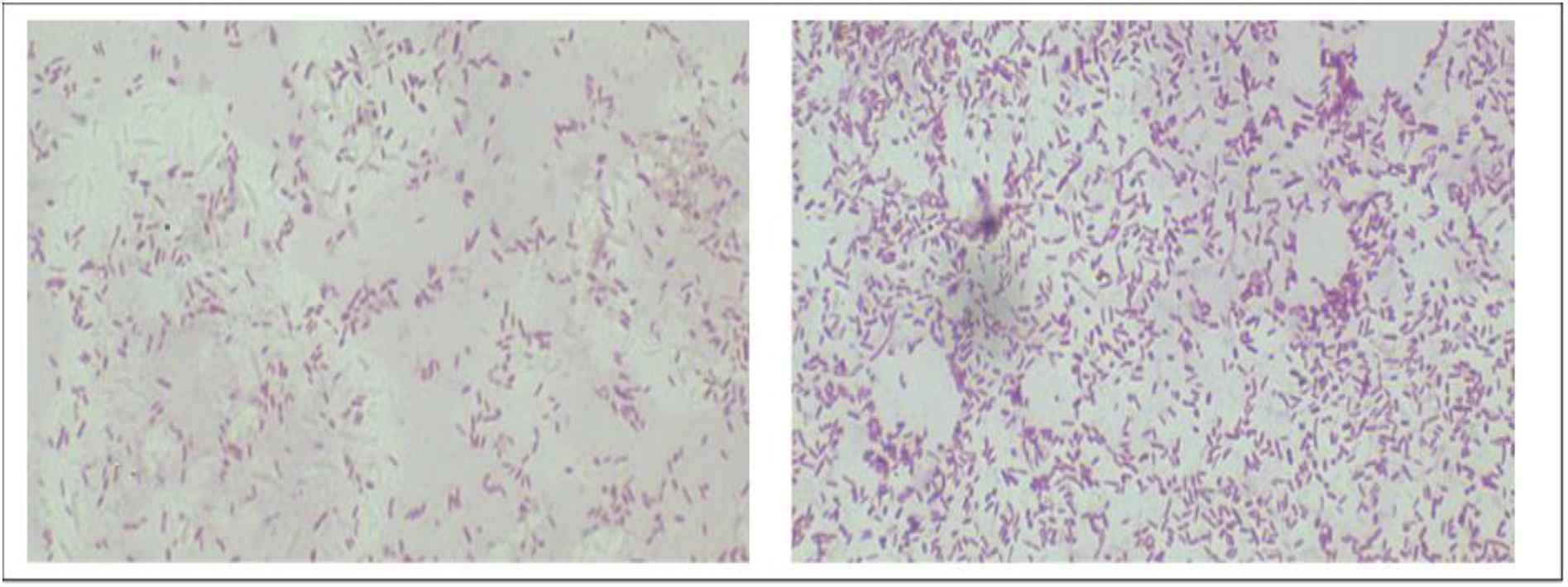
Gram staining of the MHB4CP

**Table S1.** Person’s Correlation analysis of all the considered parameters (red color indicated -1, white as 0, and blue 1)

|  | plant height | no of leaf | em diame | ophyll | mhydrate | rotein (mg/ | proline | phenol | phosphate | SFW | SDW | RFW | RDW | PH | EC | P | OC | K |
| --- | --- | --- | --- | --- | --- | --- | --- | --- | --- | --- | --- | --- | --- | --- | --- | --- | --- | --- |
| plant height | 1 |  |  |  |  |  |  |  |  |  |  |  |  |  |  |  |  |  |
| no of leaf | 0.93409 | 1 |  |  |  |  |  |  |  |  |  |  |  |  |  |  |  |  |
| pseudostem | 0.96395 | 0.9632 | 1 |  |  |  |  |  |  |  |  |  |  |  |  |  |  |  |
| chlorophyll | 0.99565 | 0.89814 | 0.94937 | 1 |  |  |  |  |  |  |  |  |  |  |  |  |  |  |
| carbohydrate | 0.85286 | 0.95806 | 0.84945 | 0.80183 | 1 |  |  |  |  |  |  |  |  |  |  |  |  |  |
| protein (mg) | 0.94689 | 0.99665 | 0.95432 | 0.91289 | 0.9675 | 1 |  |  |  |  |  |  |  |  |  |  |  |  |
| proline | 0.96334 | 0.93241 | 0.99503 | 0.95772 | 0.79692 | 0.92355 | 1 |  |  |  |  |  |  |  |  |  |  |  |
| phenol | 0.88764 | 0.91512 | 0.97638 | 0.87272 | 0.76336 | 0.88926 | 0.97611 | 1 |  |  |  |  |  |  |  |  |  |  |
| phosphate | 0.91886 | 0.95088 | 0.99073 | 0.89962 | 0.82319 | 0.93115 | 0.98387 | 0.99482 | 1 |  |  |  |  |  |  |  |  |  |
| SFW | 0.8368 | 0.93685 | 0.94955 | 0.8032 | 0.82323 | 0.90611 | 0.92976 | 0.97813 | 0.98062 | 1 |  |  |  |  |  |  |  |  |
| SDW | 0.76641 | 0.94512 | 0.85595 | 0.70632 | 0.93829 | 0.92594 | 0.80227 | 0.8433 | 0.87706 | 0.92752 | 1 |  |  |  |  |  |  |  |
| RFW | 0.85605 | 0.95317 | 0.95808 | 0.82156 | 0.84957 | 0.92603 | 0.93625 | 0.97601 | 0.9834 | 0.99874 | 0.93829 | 1 |  |  |  |  |  |  |
| RDW | 0.76848 | 0.89409 | 0.90888 | 0.73188 | 0.77224 | 0.85487 | 0.88805 | 0.96083 | 0.95521 | 0.99317 | 0.91461 | 0.9869 | 1 |  |  |  |  |  |
| PH | -0.6402 | -0.6458 | -0.4896 | -0.6054 | -0.8041 | -0.7001 | -0.4381 | -0.3036 | -0.398 | -0.3421 | -0.5604 | -0.3886 | -0.2501 | 1 |  |  |  |  |
| EC | 0.89943 | 0.98251 | 0.90138 | 0.85515 | 0.99411 | 0.9892 | 0.85767 | 0.82414 | 0.87661 | 0.86639 | 0.94149 | 0.89009 | 0.81517 | -0.7646 | 1 |  |  |  |
| P | 0.86021 | 0.96075 | 0.85531 | 0.81031 | 0.9999 | 0.9705 | 0.80399 | 0.7687 | 0.82815 | 0.82568 | 0.93651 | 0.85203 | 0.7738 | -0.8041 | 0.99531 | 1 |  |  |
| OC | 0.69489 | 0.72274 | 0.56855 | 0.65512 | 0.86587 | 0.77 | 0.51443 | 0.39607 | 0.48711 | 0.44183 | 0.65028 | 0.48599 | 0.3554 | -0.9936 | 0.83056 | 0.8657 | 1 |  |
| K | 0.45054 | 0.45131 | 0.27185 | 0.41732 | 0.65499 | 0.51432 | 0.2176 | 0.07163 | 0.17181 | 0.11829 | 0.3811 | 0.16752 | 0.0266 | -0.9721 | 0.59832 | 0.65361 | 0.94288 | 1 |

